# The prevalence of *Angiostrongylus cantonensis* in rodent hosts in China and several Asian countries: a systematic review and meta-analysis

**DOI:** 10.1101/2020.07.23.217281

**Authors:** Huifang Bai, Yizhen Cao, Yunqiu Chen, Lingmin Zhang, Chunyun Wu, Mei Cheng

## Abstract

**Background:** *Angiostrogylus cantonensis* (*A.cantonensis*) is a zoonotic parasitic nematode, with a worldwide distribution, causing eosinophilic meningitis or meningoencephalitis in humans. Although the biology of *A.cantonensis* is relatively well known, little is understood about the transmission level in different zoogeographical regions, especially in Asia. Here, to evaluate the prevalence of *A.cantonensis* in rodent hosts in China and several Asian countries, we conducted a systematic review registered with PROSPERO (CRD42020161665).

**Methods:** Records were selected systematically from 7 databases (Medline via to Pubmed, Web of science, Scopus, Google Scholar, CNKI, Wangfang, CBM). Forest plots and random-effects model were used to display pooled estimates. The Freeman-Tukey double arcsine transformation of R software was used to conduct meta-analysis and statistical significance was set at 0.05.

**Results:** A total of 67 studies met the inclusion criteria and were included in the systematic review and meta-analysis. The pooled prevalence estimates of *A.cantonensis* infection in rodents was 0.1003 (95%CI: 0.0765, 0.1268). There was significant heterogeneity in reported outcomes (p<0.0001). So we considered that there was no publication bias in the included studies.

**Conclusion:** The *A.cantonensis* infection rate among rodent hosts was still high in Asia, particularly in China, especially in *Rattus norvegicus*, and thus comprehensive measures should be taken for rodent hosts control to avoid an angiostrongyliasis outbreak. Due to the wide distribution and movement of rodent hosts, people in all regions of China, even in other Asia area live at risk of an infection. Hence, the development of more reliable diagnostic tests will be key for an effective identification of cases as well as improved patient care. Consequently, further studies are required to updated strategies for controlling *A.cantonensis* infection among human population.

## Background

*Angiostrongylus cantonensis* (*A.cantonensis*) is a zoonotic parasitic nematode, with a worldwide distribution, causing eosinophilic meningitis or meningoencephalitis in humans. Humans infection with *A.cantonensis* typically occur from intentional or unintentional ingestion of infected intermediate or paratenic hosts, sometimes due to contaminated produce.

Traditionally, with the wider domain of climate and health, the neglected tropical diseases (NTDs), are a collection of infectious diseases affected hundreds of millions of individuals living in tropical and subtropical countries. Recently, such dramatic habitat changes are linked to accelerated biodiversity loss, which has been linked to major changes in the epidemiology of human diseases: increased disease risks and the emergence of novel pathogens resulting from increased contacts among wildlife, domesticated animals, and humans[1–3]. In particular, angiostrongyliasis, as a kind of NTDs, was found in the Pacific Islands, Southeast Asia, parts of South and Central America and the Caribbean, even in European countries[4–6]. Recently, although researchers gradually pay more attention to investment towards reducing the burden of NTDs, they still collectively contribute to productively loss, illness and suffering in many countries[7,8]. Recent estimates of their overall burden suggest NTDs kill over 350,000 people per annum and cause the loss of between 27 and 56 million disability-adjusted life years[9]. To date, more than 2000 cases of eosinophilic meningitis have been recorded in China, even 100 cases in Taiwan and most human cases of angiostrongyliasis (~1300 cases) have been reported in Thailand[5,10], where between 0.3 and 2 people per 100,000 become infected annually[11,12]. These findings indicate that the extensive distribution of *A.cantonensis* in globally is high, thus putting humans as a higher risk of *A. cantonensis* infection.

In recent decades, more than 20 species of feral rodents have been categorized as pests in tropical and subtropical countries, and these species cause tremendous losses and damages to crops and food stocks[13]. Feral rodents are known to transmit diseases and act as reservoir hosts to many zoonotic parasites that pose health risks to humans, including *A.cantonensis*, hantaviruses, *Leptospira spp., Bartonella spp., Trypanosoma spp*., and *Babesia spp*.[14–16]. For *A.cantonensis*, at least 17 rodent species may behave as definitive hosts, capable of passing first stage larvae (L1) in their feces. Simultaneously, snails and slugs (intermediate hosts) ingest these larvae, which mature into the third stage larvae (L3) (infective form) that are passed on to the definitive hosts again when the definitive hosts consume infected intermediate hosts. Then, the worms travel to brain, especially in the subarachnoid space, elicits an intense eosinophilic inflammation and causes acute eosinophilic meningitis. Humans become the incidental hosts by consuming the infected intermediate or paratenic hosts (or ingestion of snail slime) [17]. Importantly, the development of control methods against zoonotic parasites depends on knowledge of their life cycles and transmission patterns in different zoogeographical regions. Therefore, it is essential to survey the relationship between parasites and rodents in order to identify the sources of zoonotic infections[18].

Surprisingly, despite rodents being the dominant vertebrate species in many of these altered tropical systems and the key hosts for numerous zoonotic microparasites (viruses, bacteria, and protists), consistent patterns between habitat fragmentation and environment change with the epidemiology of multiple parasite diseases have yet to be determined, and studies that address the context of pathogen circulation are lacking. Indeed, simultaneous to the technical and targeted approaches being, recommended by WHO are much wider attempts at sustainable development, most visible through the lens of the sustainable development goals [19]. Meanwhile, there are little diagnostic methods and effective treatment methods. Furthermore, we need do more research to solve this challenge. Here, in this study, we aim to identify the prevalence of *A.cantonensis* in rodent hosts in China and several Asian countries, providing essential recommendations for the researchers and policy makers.

## Methods

The protocol for this systematic review was defined in advance and registered with PROSPERO (CRD42020161665). This study was conducted in accordance with the PRISMA guidelines (Preferred Reporting Items for Systematic Reviews and Meta-analysis) for systematic review and meta-analysis. The PRISMA checklist (Table S1) was used to ensure inclusion of relevant information in the analysis.

### Literature search

To evaluate the prevalence of *A.cantonensis* in rodent hosts in China and several Asian countries, we carried out a systematic review of the literature (full-text) published online in the English and Chinese between May 1946 and December 2019. Records were selected systematically from 7 databases (Medline via to Pubmed, Web of science, Scopus, Google Scholar, CNKI, Wangfang, CBM). This range of years was important to study the prevalence since the first study was reported. The modified searches were performed by various combinations of the following terms using Boolean operator “AND” and “OR”: (rat lungworm OR *Angiostrongylus cantonensis* OR *Angiostrongyliasis OR Angiostrongylidae*) AND (prevalence OR distribution OR epidemic OR incidence OR frequency OR occurrence OR detection OR identification OR characterization OR investigation OR survey OR rate) AND Asia. The reference lists of selected articles also were screened manually and appropriate articles were included. Full text articles were downloaded or obtained through library resources.

### Study selection

All selected articles had to meet the following inclusion criteria: (i) a cross-sectional study; (ii) the study published between May 1946 and December 2019; (iii) the language was limited to Chinese or English; (iv) full-text article; (v) infection cases in Asia, provided geographical location; (vi) reported as animal level prevalence data, not laboratory infected animals; (vii) exact total numbers and positive cases numbers; (viii) rodent hosts numbers higher than fifty, clearly rodents species; (ix) no relevant outcomes reported.

Investigations without these criteria were excluded. Also, if the same study data were published in both English and Chinese sources, the articles with less detailed information would be excluded from this study. When any authors found articles difficult to judge, the first author was consulted and eventual differences discussed to reach a consensus decision on whether to include or exclude the article.

### Quality assessment

The included articles were evaluated using a 10-item quality appraisal checklist (Text S1). An item would be scored ‘0’ if it was answered ‘no data available’ or ‘unclear’; if it was answered ‘yes’, then the item scored ‘1’. Article quality was assessed as follows: low quality = 0–3; moderate quality = 4–7; high quality = 8–10.

### Data extraction

For the data extraction, the two independent assessors applied the inclusion criteria selected the articles and extracted the data. Discrepancies were resolved by consensus. The detailed characteristics of each study were extracted using a pre-designed data-collection excel form. Information was collected as followings: first author, year of publication, location, geographical region, sample size, number of positive cases, diagnostic methods, types of rodent hosts, quality score of each study.

### Statistical analysis

The statistical software used in the analysis was R software version (3.6.3). Estimated pooled prevalence and 95% confidence intervals (CI) were calculated with the Freeman-Tukey double arcsine transformation. Heterogeneity testing was performed using the I^2^ and Cochran’ Q statistic methods. A significant value (p<0.05) in the analysis suggested a real effect difference. The I^2^ values of 25%, 50% and 75% were considered as low, moderate and high heterogeneity, respectively. The risk of study publication bias was assessed using the funnel plots, and the Egger’s regression test.

Also a significant value (p<0.05) in the analysis suggested a real effect difference. Furthermore, subgroup analysis were done if the heterogeneity were overlapping moderate I^2^. Four potential sources of heterogeneity were examined: country, animal species, publication year, quality score. The Q and I statistics values were calculated for each subgroup to determine the effective factors on the prevalence *A.cantonensis* and heterogeneity about all the included studies.

## Results

### Study search and included studies

A total of 2103 articles were screened in the initial and 67 were obtained for the final analysis. Fig 1 showed our systematic workflow for identifying, screening and including studies in this study. Hence, 59 articles from China, 1 from Japan, 4 from Thailand, 1 from India and 2 from Philippines. The complete list of included articles with the extracted data can be found in the Fig 2 and Supporting information (Text S2). A total of 45196 rodents are dissected and 4954 are identified with *A.cantonensis* infection. The detailed characteristics of each study were provided in Table 1.

**Fig 1.**
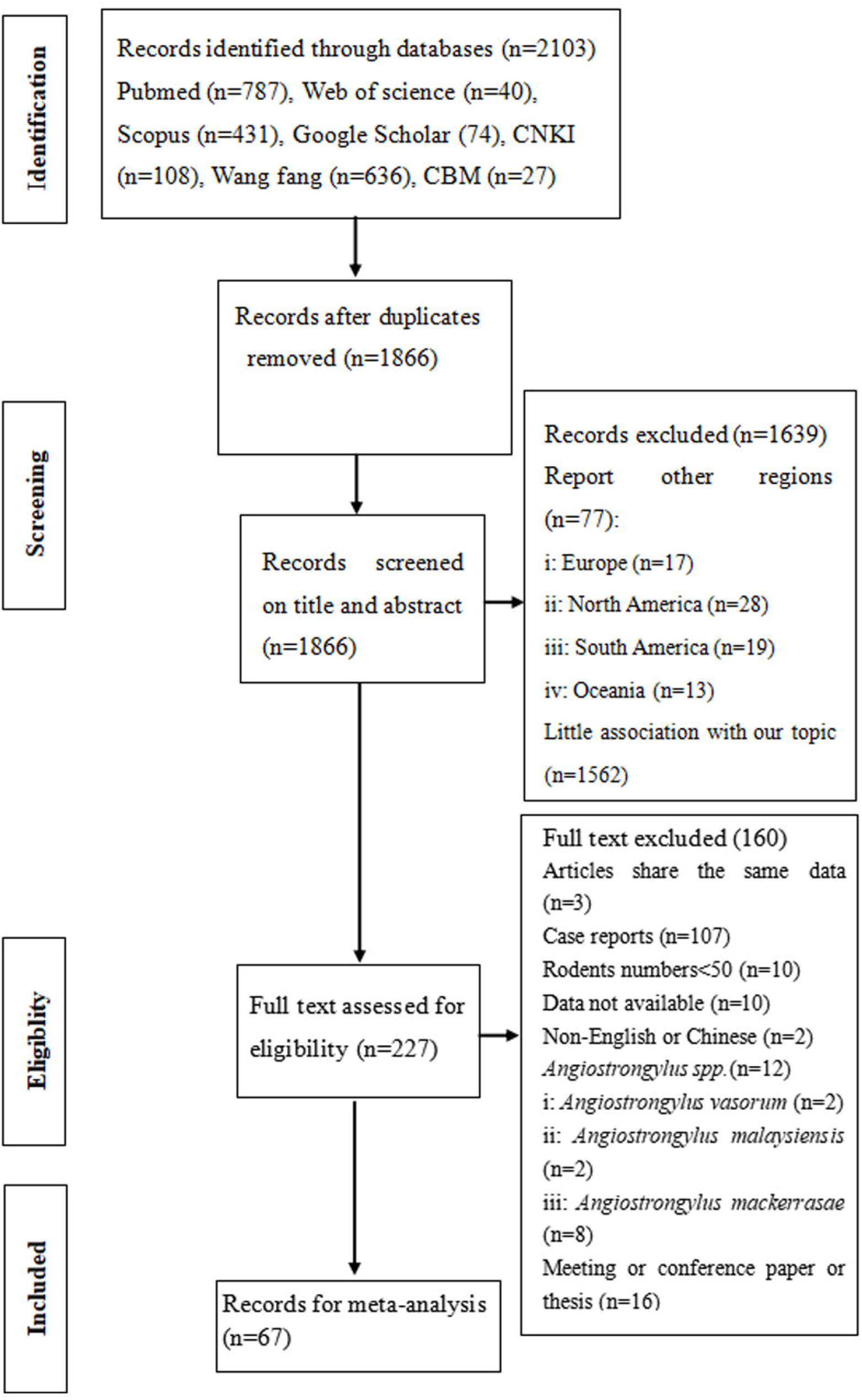
Flow-chart of systematic review of *A. cantonensis*

**Fig 2.**
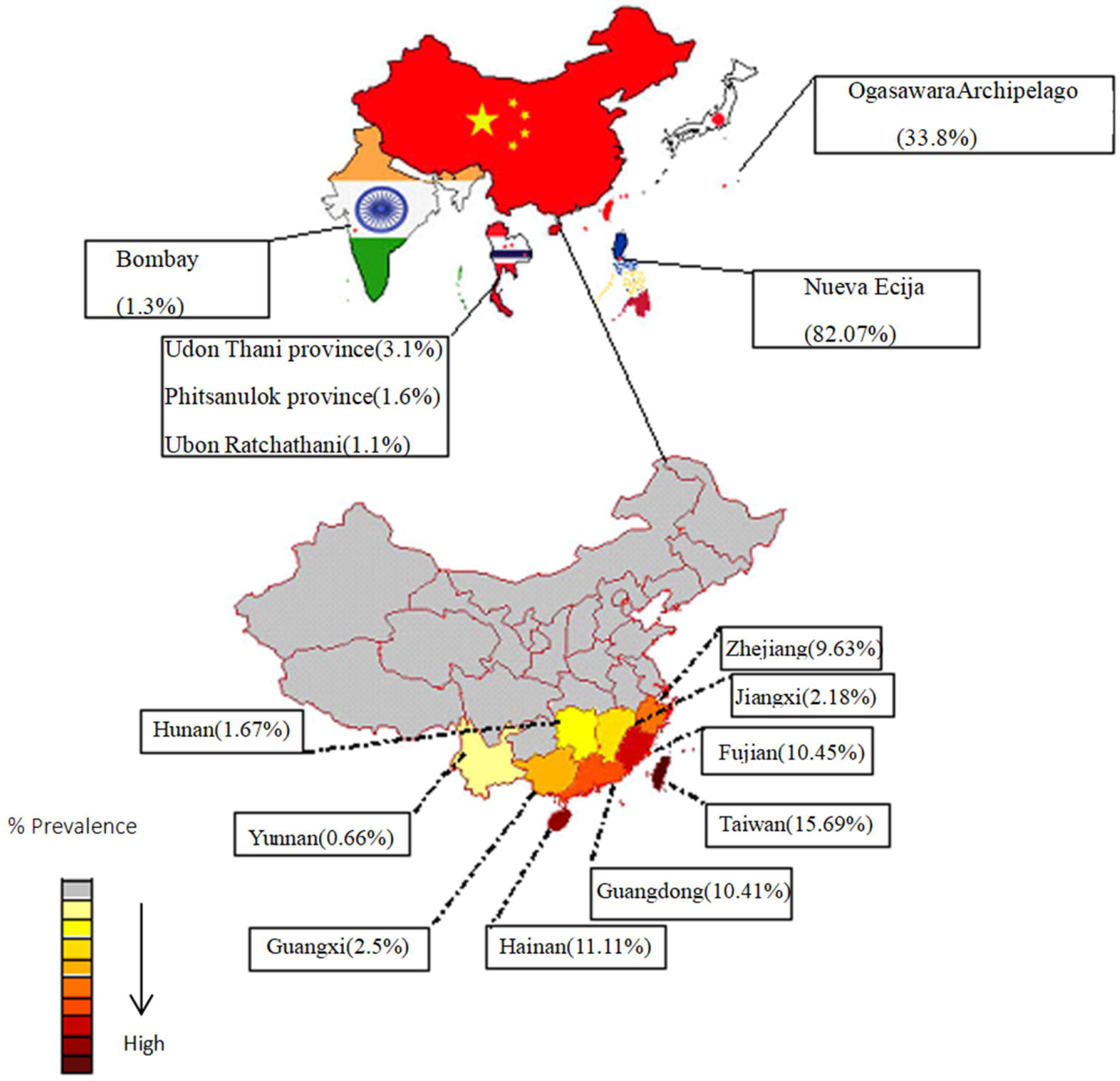
Map of the geographical distribution in Asian countries of the surveys

**Table 1.**
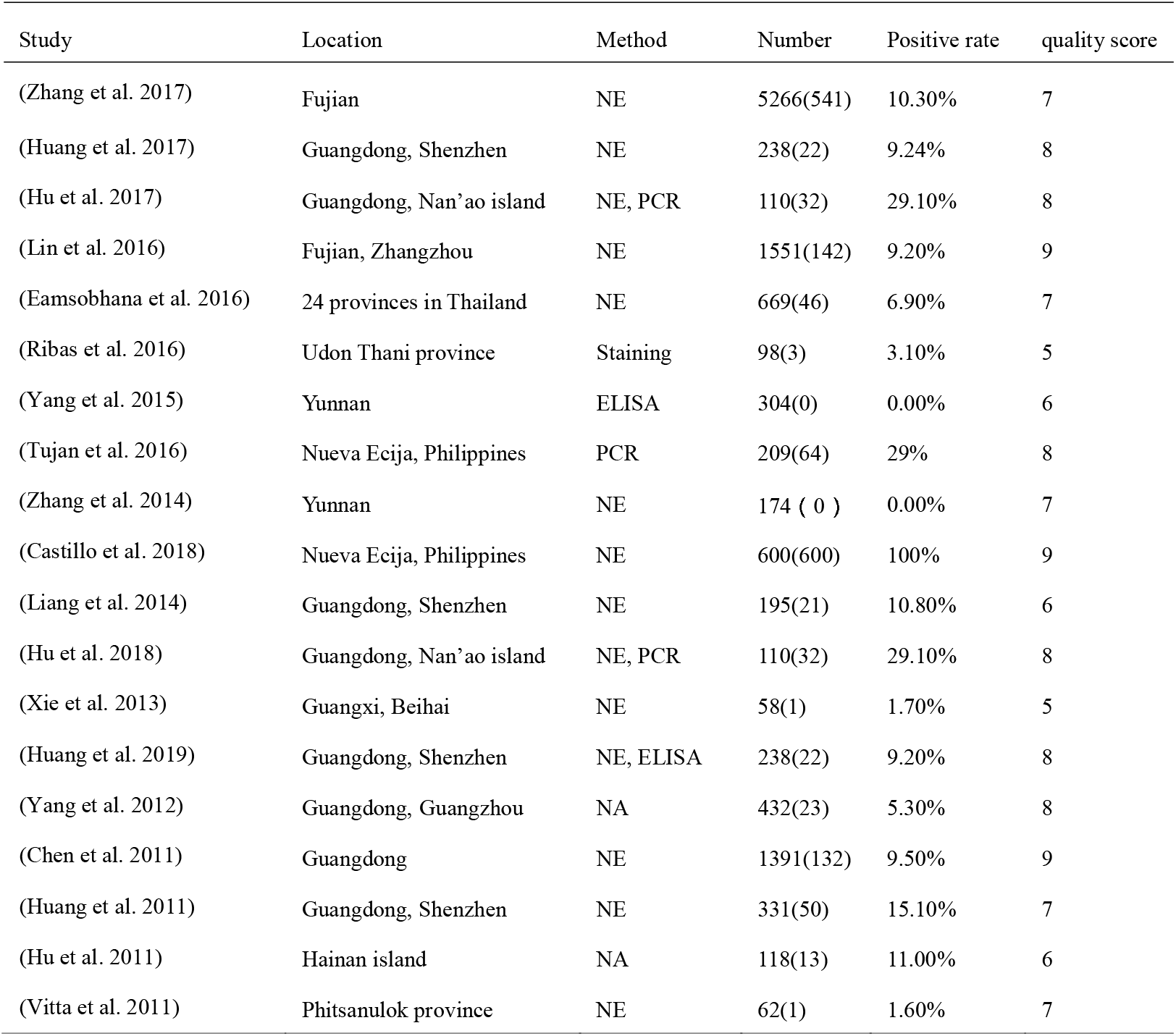
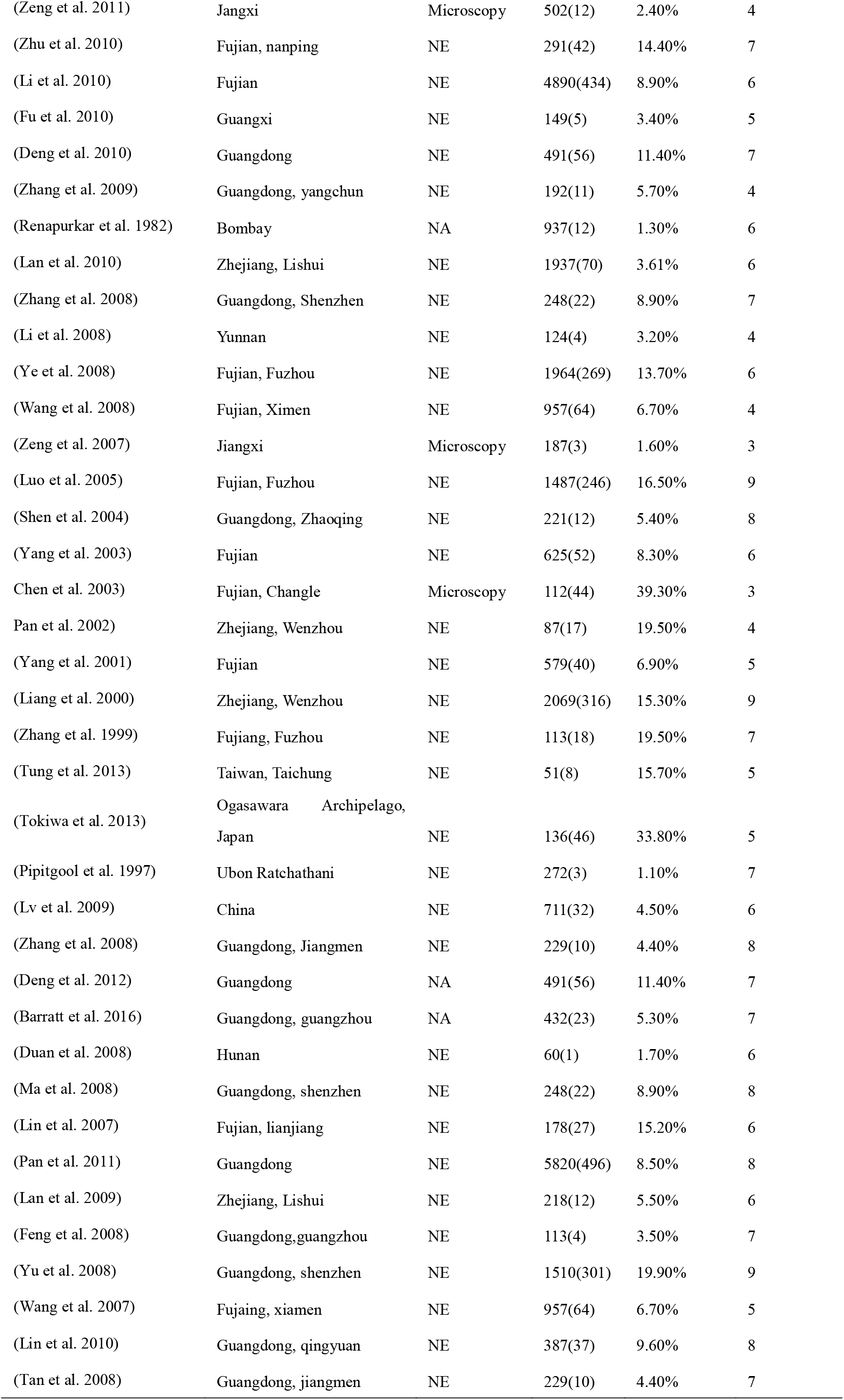
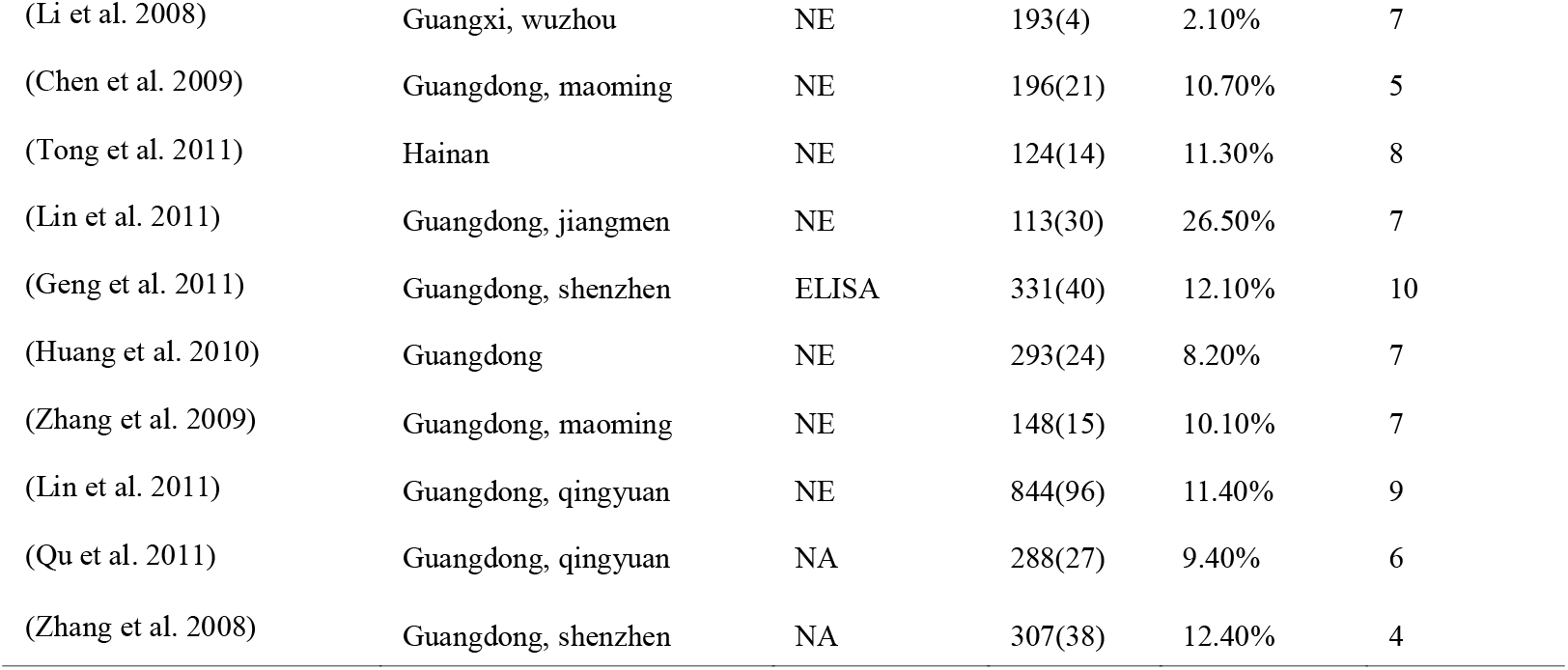
Characterizatics of the eligible studies

### Pooling and heterogeneity analysis

The pooled prevalence estimates of *A.cantonensis* infection in rodent hosts with individual studies are shown in a forest plot (Fig 3). A substantial heterogeneity is observed among studies (χ^2^ = 4979.77, P<0.05; I^2^=99.0%). When calculated using a random-effects model, the overall infection rate is 0.1003 (95%CI: 0.0765,0.1268).

**Fig. 3.**
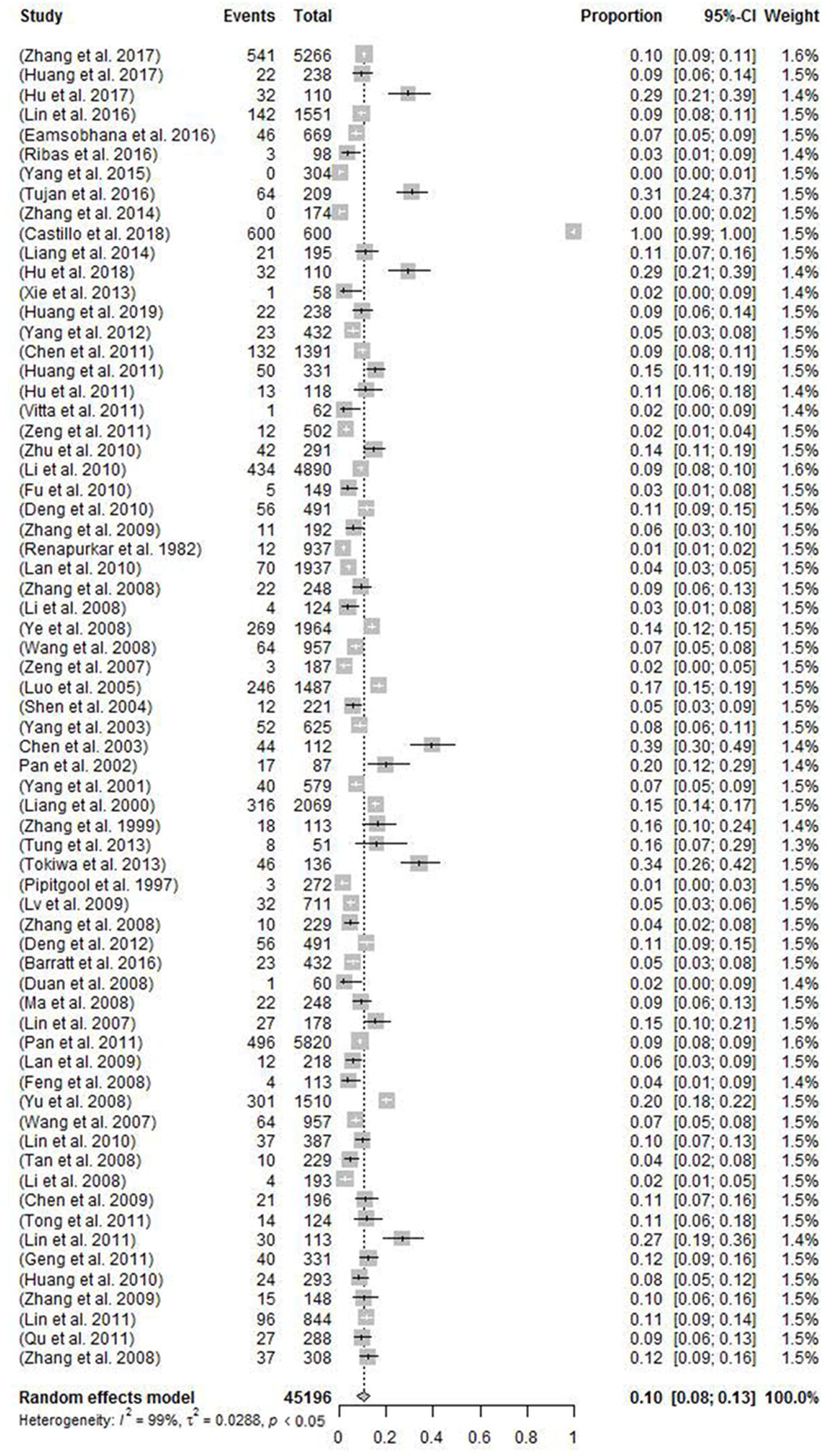
Random-effects model meta-analysis of *A.cantonensis* infection in Asian countries

The estimates of infection rates for different subgroups and heterogeneities are presented in Table 2 and Fig 4. All pooled infection rates for each subgroup are calculated using a random-effects model because of the observed high heterogeneity among studies within subgroups. Base on study periods, the estimate is significantly increased since 2008 (≤2008: 0.0834 (95%CI: 0.0595, 0.1107)vs > 2008: 0.111 (95%CI: 0.0758, 0.1518)). In terms of the rodent hosts species, the estimate is highest in *Rattus norvegicus* (0.1441, 95%CI: 0.1100, 0.1818), followed by *Rattus flavipectus* (0.0428, 95%CI: 0.0062, 0.0987), *Bandicota indica* (0.0409, 95%CI: 0.0000, 0.1240), *Mus musculus* (0.0101, 95%CI: 0.0041, 0.0187), *Rattus rattus* (0.0093, 95%CI: 0.0004, 0.0299), *Rattus losea* (0.0044, 95%CI: 0.0000, 0.0176), and the lowest is *Suncus Murinus* (0.0006, 95%CI: 0.0000, 0.0086). In terms of Asian region, China (0.0893, 95%CI: 0.0773, 0.1020) is higher than Thailand (0.0299, 95%CI: 0.0053, 0.0700). Finally, in terms of score, the estimate is highest in high score(0.1266, 95%CI: 0.1021, 0.1560), followed by low score (0.0892, 95%CI: 0.0420, 0.1509), the lowest was middle score (0.0728, 95%CI: 0.0583, 0.0888).

**Table.2.** Subgroup meta-analysis of *A.cantonensis* in Asian countries

| variable | population | No.of<br>infected<br>rodents | No. of<br>total<br>rodents | Pooled<br>estimate<br>Prevalence<br>(%) | 95%CI | Heterogeneity(Q) |  |  | Egger'test |  |
| --- | --- | --- | --- | --- | --- | --- | --- | --- | --- | --- |
|  |  |  |  |  |  | Q | I<sup>2</sup>(%) | P-value | t | P |
| **Asia region** |  |  |  |  |  |  |  |  |  |  |
| **China** | 42213(4179) | 4179 | 42213 | 0.0893 | (0.0773,0.1020) | 1005.38 | 95.1 | <0.0001 | -0.5678 | 0.5724 |
| **Thailand** | 1101(53) | 53 | 1101 | 0.0299 | (0.0053,0.0700) | 19.67 | 93.9 | 0.0002 | -1.0502 | 0.4038 |
| **Japan** | 136(46) | 46 | 136 |  |  |  |  |  |  |  |
| **Philippines** | 809(664) | 664 | 809 |  |  |  |  |  |  |  |
| **india** | 937(12) | 12 | 937 |  |  |  |  |  |  |  |
| **Score** |  |  |  |  |  |  |  |  |  |  |
| **≤4 (low)** | 2469(192) | 192 | 2469 | 0.0892 | ( 0.0420,0.1509 ) | 139 | 96.8 | <0.0001 | 1.0588 | 0.3304 |
| **> 4 to ≤7**<br>**(middle)** | 24578(2103) | 2103 | 24578 | 0.0728 | ( 0.0583,0.0888 ) | 674.28 | 95.4 | <0.0001 | -0.8451 | 0.4035 |
| **> 7 to ≤10**<br>**(high)** | 18149(2659) | 2659 | 18149 | 0.1266 | (0.1021,0.1560) | 421.26 | 96.5 | <0.0001 | 0.4219 | 0.6781 |
| **Host species** |  |  |  |  |  |  |  |  |  |  |
| ***Rattus***<br>***norvegicus*** | 18479(3048) | 3048 | 18479 | 0.1441 | (0.1100,0.1818) | 2320.32 | 98 | <0.0001 | -0.4383 | 0.663 |
| ***Rattus***<br>***flavipectus*** | 7840 ( 918 ) | 918 | 7840 | 0.0428 | (0.0062,0.0987) | 2230.08 | 98.3 | <0.0001 | -0.409 | 0.6846 |
| ***Suncus***<br>***Murinus*** | 5758 ( 179 ) | 179 | 5758 | 0.0006 | ( 0.0000,0.0086 ) | 164.89 | 87.2 | <0.0001 | -0.7805 | 0.4416 |
| ***Mus musculus*** | 1137 ( 22 ) | 22 | 1137 | 0.0101 | ( 0.0041,0.0187 ) | 34.88 | 44.6 | 0.2469 | -2.2943 | 0.0292 |
| ***Rattus losea*** | 1697 ( 69 ) | 69 | 1697 | 0.0044 | ( 0.0000,0.0176 ) | 25.5 | 62.3 | 0.0841 | 1.8715 | 0.0797 |
| ***Bandicota***<br>***indica*** | 800 ( 66 ) | 66 | 800 | 0.0409 | ( 0.0000,0.1240 ) | 90.17 | 93.1 | <0.0001 | 0.3826 | 0.7109 |
| *Rattus rattus* | 797(19) | 19 | 797 | 0.0093 | ( 0.0004,0.0299 ) | 35.34 | 79.3 | 0.0008 | 0.8389 | 0.4179 |
| other species | 4269(415) | 415 | 4269 |  |  |  |  |  |  |  |
| **publication year** |  |  |  |  |  |  |  |  |  |  |
| ≤2008 | 14007(1602) | 1602 | 14007 | 0.0834 | (0.0595,0.1107) | 665.2 | 97.1 | <0.0001 | -1.833 | 0.0798 |
| > 2008 | 31189(3352) | 3352 | 31189 | 0.111 | (0.0758,0.1518) | 4311.74 | 99.1 | <0.0001 | 0.6216 | 0.5377 |
| **Overall** | **49156(4954)** | **4954** | **49156** | **0.1003** | **(0.0765,0.1268)** | **4979.77** | **99.0** | **<0.0001** | **0.1075** | **0.9147** |

**Fig. 4.**
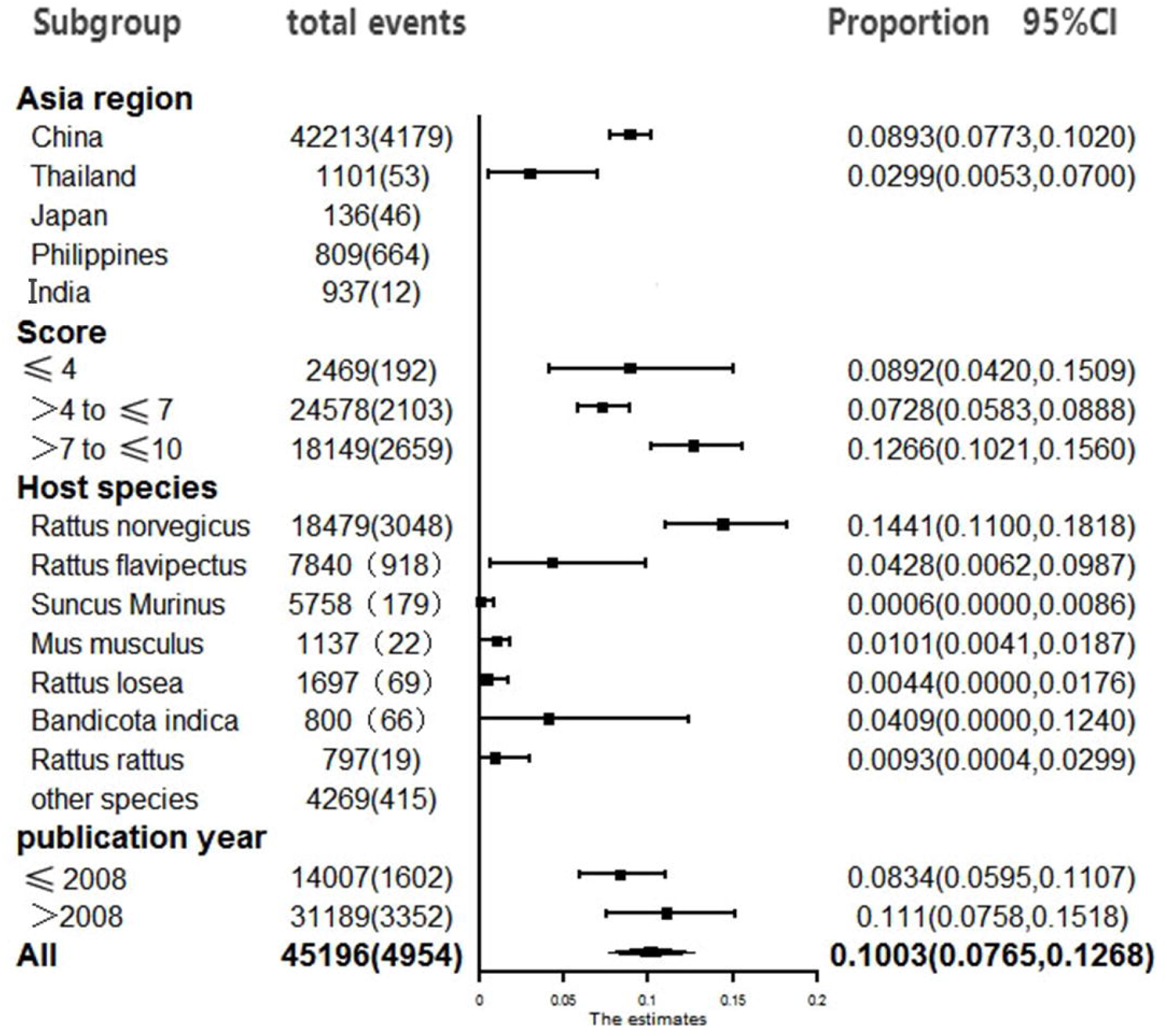
Subgroup forest plot of infection *A. cantonensis* in Asian countries

### Publication bias

Despite the significant heterogeneity, the funnel plot displayed a symmetric spread of studies in terms of relative weight and effect size, thereby we could not directly judge whether there was publication bias in the included studies (Fig. 5). The results of Egger’s test showed P=0.9147 (P>0.05). Therefore, we considered that there was no publication bias in the included studies. Notably, the total number of studies was reasonable, and the individual studies were of variable sample size. Moreover, Duval and Tweedie’s trim and fill procedure for the detection of publication bias did not support the possibility of missing studies from the analysis.

**Fig. 5.**
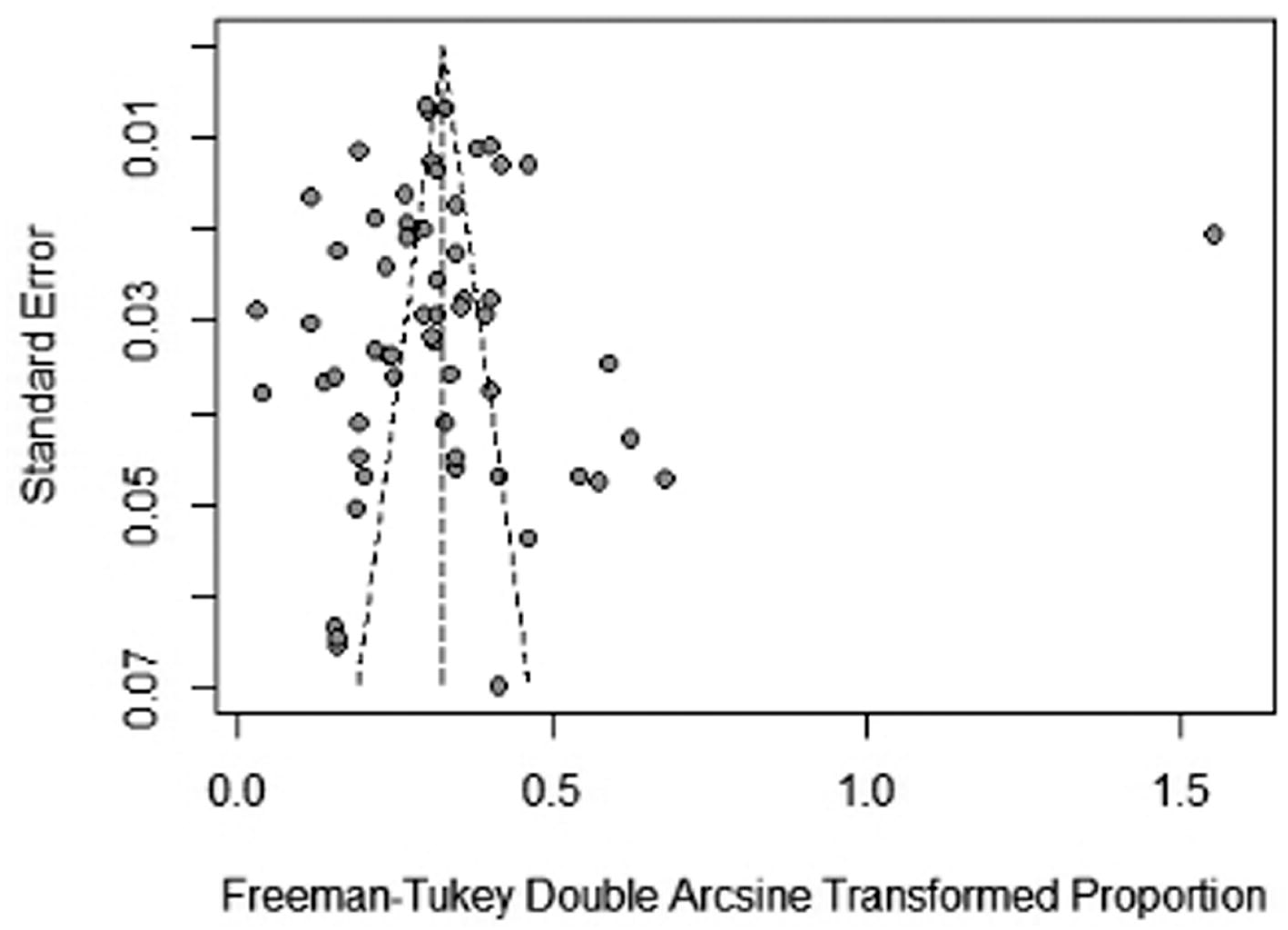
Funnel plot of the arcsine transformed infection rates of *A.cantonensis* in Asian countries

## Discussion

In previous study, rodents may play an important role in the transmission of certain zoonotic parasites to humans[18]. For *A.cantonensis*, the estimate is highest in *Rattus norvegicus, Rattus flavipectus* and *Bandicota indica*, followed by *Mus musculus, Rattus rattus*, *Rattus losea*, and *Suncus Murinus*. The difference of the infection rates among seven kinds of rats may be relative to the factors: the susceptibility to *A. cantonensis*, rat’s habitat, activity range and hunted objects, and so on. For example, the front several rodents(*Rattus norvegicus, Rattus flavipectus* and *Bandicota indica*) : (i) are considered as dominant species in several Asian countries[20,21]; (ii) live close to human dwellings [22]; (iii) they were also known to prey on snails not plants [23]. This result is consistent with Chen D’s report[24].

Before 2008, there were outbreaks in Beijing, Wenzhou, Guangzhou and other areas in China, particularly, 160 cases infected in Beijing[25]. Then, investigators completed additional research across China by the national angiostrongyliasis control programme. Prevalence of *A. cantonensis* in different countries of the world is most likely related to several issues such as climate changes and land alterations, changes in the behavioral and physiological patterns and species compositions of hosts, climate changes affect behavior of hosts having a key role in spread of *A.cantonensis*, as well, lead to variation in seasonal / temperature dependent activities in the snail and rodents. These changes affect the frequency of transmission of parasites and severity of infection. The environment change is including soil, water and air. Soil is the upper covering layer of the earth, consisting of three subphases (water, gas and solids) which combine to describe the overall mechanics and other properties. Soil is essential to the natural history of many parasites and their vectors; examples include eggs of geohelminths[26], larvae of Tsetse flies[27] and burrows of mammals fed on by triatominae insects[28]. With the land surface temperatures increasing, the chemical elements and physical properties of the soil have changed. All these soil changes can lead the definitive host survival rate up or down. Also, water is important to the natural history of many parasites or their vectors equally. Thermal tolerance may be a critical issue for many water-based, or semiaquatic organisms involved in the life cycles-including insects, freshwater snails, fish, crabs, copepods, crayfish and insects. Poikilothermic ectotherms such as these consume oxygen based on the water or temperature until some threshold temperature where ATP supply and demand is over whelmingly disrupted and the organism dies. The life cycles of several NTDs including Schistosoma, the food-borne trematodes and *A.cantonensis* involve intermediate hosts that may inhabit and reproduce in water bodies with thermal stratification, such as lakes. Analysis of historic data indicates that global warming is associated with changes to lake stratification that are dependent on lake morphometry[29]. How the intermediate hosts will respond over the coming decades is unclear, but evidence suggests evolution may have led to divergent populations of copepods that have adapted to warmer or colder conditions. The central tenet of climate change is the forcing effects of so-called ‘greenhouse’ gasses including CO_2_ and aerosols. The relationship between surface air temperature (as predicted by climate projections) is generally assumed to be correlated, over decadal scales, with the ground surface temperature, but over shorter time scales there may be considerable variability. Abiotic changes to the soil as a result to changes in air temperature may affect the natural history of a wide range of NTDs as diverse as trypanosomes cestodes and nematodes. In such environments, more case will be induced, a high prevalence of parasitic infections (hantavirus, Yersinia pestis) is often found in various rodent species[30]. In normal, *Rattus spp*. feces are often found in water or soil. The major intermediate hosts (e.g. *Pomacea canaliculata* and *Achatina fulica*)[31] maybe infected by ingesting feces. Typically, soil and water properties’ changes will affect the intermediate hosts and definitive hosts living time. Their living stages are an essential component of the *A.cantonensis* infection life cycle. Moreover, with the wide habitat alternation, the hosts may adapted and mitigated quickly to modify emissions and habitats. Finally, all factors mentioned earlier leads cases increased year by year.

## Limitations

Though this study provided information for the prevalence of *A.cantonensis* in rodent hosts in Asia, several limitations may merit attention. First, the estimations of infection rates using a random-effect model was unavoidable, then subgroup analysis was employed but could not fully explain the source of heterogeneity. Secondly, our study didn’t include all Asian countries because the relevant data were not available or incomplete. Finally, in this study, we only focused on *Rattus norvegicus, Rattus flavipectus, Bandicota indica, Mus musculus, Rattus rattus, Rattus losea* and *Suncus Murinus*, because they were common and most important rodent hosts. However, other rodents, like *Rattus tiomanicus, Rattus exulans Bandicota savilei* and *Maxomys surifer*, though diffiiculty to collect them but should not be neglected. To fully elucidate the prevalence of angiostrongyliasis, further studies should pay attention to the other species rodent hosts.

## Conclusion

In conclusion, our systematic review and meta-analysis estimated a pool national prevalence of *A.cantonensis* in rodent hosts in Asian areas, particularly in south China, which provided a scientific basis for prevention a control of the spread of angiostrongyliasis. The *A.cantonensis* infection rate among rodent hosts was still high in China, especially in *Rattus norvegicus*, and thus comprehensive measures should be taken for rodent host control to avoid a potential angiostrongyliasis outbreak. Moreover, public health officials, epidemiologists, researchers, clinical technicians, medical practitioners, parasitologists, and veterinarians, as well as the general public, should be aware of such risks, and integrated strategies should be taken to reduce or eliminate such risks[32]. Consequently, further studies are required to updated strategies for controlling *A.cantonensis* infection risk among human population.

## Supplementary information

**Additional fle 1: Table S1.** PRISMA checklist. **Text S1.** Quality assessment checklist. **Text S2.** References of the selected articles included in the systematic review.

## Abbreviations

*A.cantonensis*: *Angiostrongylus cantonensis*
CI: Confidence interval
NTDs: the neglected tropical diseases
L1: first stage larvae
L3: third stage larvae
WHO: World Health Organization
PRISMA: Preferred Reporting Items for Systematic Reviews and Meta-analysis
ATP: Adenylate triphosphate.

## Acknowledgements

Not applicable.

## Author Contributions

Conceptualization: BHF, CM.

Data curation: BHF, CYZ, CYQ.

Formal analysis: BHF, CYZ, CYQ, ZLM.

Investigation: CYQ, ZLM, WCY.

Methodology: BHF, CM.

Validation: ZLM, CM.

Visualization: CYZ, CYQ, ZLM, WCY.

Writing-original draft: BHF, CYZ.

Writing-review & editing: BHF, CYZ, CYQ, CM.

## Availability of data and materials

All data are included in this article.

## Ethical approval and consent to participate

Not applicable.

## Consent for publication

Not applicable.

