## Supplement Text S1 for "The prevalence of *Angiostrongylus cantonensis* in rodent hosts in China and several Asian countries: a systematic review and meta-analysis"

Text S1: Quality assessment checklist

The following items were examined and given a score based on a simple scale system (1 for ''yes'', 0 for ''no data avaliable'' or “unclear”).

1. Was the research objective clearly stated?
2. Was the sampling area clearly described with reference to the location?
3. Was the period of the study stated?
4. Was the target sample a close representation of the general population?
5. Was some form of random selection used to select the samples
6. Was a justified and satisfactory sample size calculated?
7. Were the sample processing and diagnostic method clearly described?
8. Were the subjects categorised by sex or age?
9. Was the sample type mentioned?
10. Was the study have a clear method to assess the outcome?
